# Decoding Liver Cancer Prognosis: From Multi-omics Subtypes, Prognostic Models to Single Cell Validation

**DOI:** 10.1101/2024.11.04.610003

**Authors:** Yanbin Wang, Yuqi Wu, Hong Zhang, Xinyue Liu, Jing Ling, Xiao Zhou, Anping Song, Li Sun, Hong Qiu, Xianglin Yuan, Hua Xiong, Yanmei Zou

## Abstract

**Purpose:** Hepatocellular carcinoma (HCC) is a highly aggressive tumor characterized by significant heterogeneity and invasiveness, leading to a lack of precise individualized treatment strategies and poor patient outcomes. This necessitates the urgent development of accurate patient stratification methods and targeted therapies based on distinct tumor characteristics.

**Experimental Design:** By integrating gene expression data from The Cancer Genome Atlas (TCGA), International Cancer Genome Consortium (ICGC), and Gene Expression Omnibus (GEO), we identified subtypes through a multi-omics consensus clustering approach amalgamated from 10 clustering techniques. Subsequently, we developed a prognostic model, employing machine learning algorithms, based on subtype classification features. Finally, by analyzing single cell sequencing data, we investigated the mechanisms driving prognostic variations among distinct subtypes.

**Results:** First, we developed a novel consensus clustering method that categorizes liver cancer patients into two subtypes, CS1 and CS2. Second, we constructed a prognostic prediction model, which demonstrated superior predictive accuracy compared to several models published in the past five years. Finally, we observed differences between CS1 and CS2 in various metabolic pathways, biological processes, and signaling pathways, such as fatty acid metabolism, hypoxia levels, PI3K-AKT and MIF signaling pathway.

## Introduction

Liver cancer is currently one of the most common tumors, and among primary liver cancers, hepatocellular carcinoma (HCC) is the most common pathological type. In 2022, liver cancer saw widespread new cases globally, and its incidence rate has been gradually increasing in recent years, ranking 6th among all tumors worldwide^[1]^. Meanwhile, liver cancer ranks 3rd among tumors causing patient deaths, with a relative 5-year survival rate of about 18%, highlighting the significantly poor prognosis of this disease^[2]^.

With the advancement of current treatment technologies, there are ongoing attempts to employ advanced treatment methods for innovative liver cancer therapy. Nevertheless, up to now, the progress of strategies aimed at treating liver cancer, especially unresectable liver cancer, has been limited. For instance, targeted therapies and immunotherapies have been the focus of extensive research efforts and have shown promising results in some patients. However, a significant number of patients still do not obtain clinical benefits from these approaches^[3]^. This may be due to the significant heterogeneity of liver cancer; classifying liver cancer patients according to molecular subtypes might help overcome this difficulty^[4]^. This urgently necessitates leveraging large-scale multi-omics data and advanced machine learning algorithms to identify biomarkers capable of efficiently predicting clinical treatment outcomes for liver cancer patients.

In this study, we combined mRNA, long non-coding RNA (lncRNA), and microRNA (miRNA) expression profiles, genomic mutations, and epigenomic DNA methylation data, using a consensus strategy integrating 10 multi-omics clustering methods to develop comprehensive consensus subtypes of liver cancer, and conducted preliminary exploration of the molecular characteristics and potential clinical applications of the two subtypes. Subsequently, we further expanded the clinical application value of the subtype. Based on the gene expression profiles of two subtypes, we used a combination of 101 machine learning algorithms to construct the best-performing prognostic model. This prognostic model showed excellent prognostic value in both training, internal and external validation sets, and outperformed several other prognostic models in comparisons. Finally, we validated the subtypes at the single-cell sequencing cell level using the same typing method, and further explored the possible causes and mechanisms of prognostic differences in patients with different subtypes.

## Materials and Methods

### Data collection and processing

Download the molecular profiles of the TCGA-LIHC cohort dataset from the TCGA database, including 339 LIHC cases with complete transcriptome expression, somatic mutations, copy number alterations (CNA), DNA methylation, and clinical information. The transcriptome profiles of mRNA and lncRNA were obtained using the “TCGAbiolinks” R package^[5]^, and GENCODE27 was used to convert Ensemble IDs to gene symbols; Somatic mutation data were also obtained using “TCGAbiolinks” R package and processed with the “maftools” R package^[6]^; DNA methylation profiles and clinical information were downloaded from UCSC Xena (https://xenabrowser.net/); Meanwhile, we also downloaded 445 LIHC cases with complete transcriptome expression and clinical information from the LIHC.JP (https://dcc.icgc.org/) and GSE14520 (https://www.ncbi.nlm.nih.gov/geo/, accession number GSE14520) cohorts; The single-cell sequencing datasets are sourced from the GEO database (https://www.ncbi.nlm.nih.gov/geo/, accession number GSE229772).

### Construct multi-omics consensus clustering classification and perform validation

In this study, we utilized the “getElites” function from the multi-omics integration and visualization (MOVICS) package to screen gene features in cancer subtypes^[7]^. For continuous variables (mRNA, lncRNA, miRNA, and methylation), we set the “method” parameter of the “getElites” function to “mad” to screen the top 1000 genes with the greatest variability; For binary variable gene mutation data, we conducted screening based on mutation frequency by setting the “method” parameter to “freq” to identify genes with a mutation frequency exceeding 10. The results from these five dimensions were incorporated into our study for further analysis. After the initial feature selection, we used the “getClustNum” function in the MOVICS package to further determine the optimal number of clusters for our study and ultimately decided to divide them into two subtypes. Next, we applied the “getMOIC” function for clustering analysis. We used 10 clustering algorithms (CIMLR, ConsensusClustering, SNF, iClusterBayes, PINSPlus, moCluster, NEMO, IntNMF, COCA, and LRA) as input for the “methods.list” parameter and employed the default parameters provided by the MOVICS package. As a result, we obtained the clustering results for each method. After calculating the clustering results of the 10 methods, we integrated the results of different algorithms based on the concept of consensus clustering using the “getConsensusMOIC” function, obtaining the final clustering results. Subsequently, we conducted nearest template prediction(NTP) analysis, performed subtyping in the external ICGC-JP cohort, and validated the subtyping method.

### Molecular Characteristics of Multi-Omics Consensus Clustering Subtypes

We analyzed the mutation status using the “maftools” R package, then used the limma method to analyze the differentially expressed genes between the two subtypes^[8]^, and performed enrichment analysis with the clusterProfiler R package^[9]^.

### Cell, signaling pathways, stemness, and other phenotypic characteristics of multi-omics consensus clustering subtypes

Using ssGSEA analysis, we calculated the enrichment scores of different immune cell characteristics between the two subtypes based on immune cell gene sets reported in the literature^[10]^; Next, we analyzed the activation status of cancer pathways and some biological processes in patients using the GSEA algorithm, and analyzed the sensitivity of the two subtypes to conventional therapeutic drugs. Both of these steps were performed using the “GSVA” R package^[11]^. The expression data from CCLE were sourced from the Broad Institute Cancer Cell Line Encyclopedia (https://sites.broadinstitute.org/ccle/); The CTRP v.2.0 (https://portals.broadinstitute.org/ctrp) and PRISM dataset (24Q2; https://depmap.org/portal/prism/) were utilized to acquire drug sensitivity data; The half-maximal inhibitory concentration (IC50) was used as an indicator of drug sensitivity.

### Constructing a prognostic prediction model based on the consensus clustering of multi-omics features

To build the prognostic model, we integrated 10 machine learning algorithms and 101 algorithm combinations^[12]^. The integrative algorithms included random survival forest (RSF), elastic network (Enet), Lasso, Ridge, stepwise Cox, CoxBoost, partial least squares regression for Cox (plsRcox), supervised principal components (SuperPC), generalised boosted regression modelling (GBM), and survival support vector machine (survival-SVM). The signature generation procedure was as follows: (a) Univariate Cox regression identified prognostic genes in the TCGA-LIHC; (b) Then, 101 algorithm combinations were performed on the prognostic genes to fit prediction models based on the leave-one-out cross-validation (LOOCV) framework in the TCGA-LIHC cohort and incorporated 10 machine learning algorithms, including CoxBoost, stepwise Cox, Lasso, Ridge, Enet, survival-SVMs, GBMs, SuperPC, plsRcox, and RSF, creating a total of 101 different combinations of these algorithms. Ultimately, these 101 algorithm combinations will be utilized to develop the most predictive prognostic model with the best C-index performance. We then scored each sample in the training and validation sets based on the model and classified the samples into high-risk and low-risk groups according to the optimal cut-off value of scores, and the “survminer” R package (https://github.com/kassambara/survminer/tree/master) was used to determine the optimal cut-off value. Cox regression and Kaplan–Meier analyses were performed via the “survival” R package (https://CRAN.R-project.org/package=survival).

### Comparison of Prognostic Prediction Models

We evaluated the prognostic predictive ability of all models included in the study using the C-index in each cohort. To enhance the clinical utility of the model, we constructed a nomogram based on factors obtained from multivariable Cox regression. We plotted time-dependent C-index curves and calibration curves to describe accuracy and used decision curve analysis to calculate the clinical benefit for patients.

### scRNA-seq Data Processing

We utilized the “Seurat” R package to process single-cell sequencing data from the GSE229772 dataset^[13]^. We removed samples with over 10% mitochondrial genes and over 5% ribosomal genes, and filtered out genes expressed in fewer than 3 cells. Next, we identified the most variable genes using the FindVariableFeatures function from the Seurat R package, conducted dimensionality reduction via principal component analysis (PCA), and identified a total of 13 clusters through t-SNE clustering analysis. Finally, we annotated the cell populations by combining the expression of characteristic genes in each cell cluster with the expression of commonly used cell-type-specific genes across the clusters.

### Characteristic Analysis of Different Cell Subpopulations

We used the “irGSEA” R package^[14]^, employing 6 methods (AUCell, UCell, singscore, ssGSEA, JASMINE, and viper) for pathway enrichment analysis among different cell populations, and performed a comprehensive evaluation of the differential analysis results using robust rank aggregation (RRA) to identify gene sets significantly enriched in most enrichment analysis methods. Moreover, we additionally utilized the “scMetabolism” R package to compare the metabolic difference across various cell types^[15]^. The metabolic activity of each cell was quantified through integrating the gene sets contained in the KEGG database. Subsequently, we used the “CellChat” R package to explore the interaction patterns among cell populations^[16]^, defining ligands and receptors as outgoing and incoming signals, respectively.

### Characteristics of Different Subtypes of Malignant Cells at the Single-Cell Level

We applied the “CopyKAT” R package to analyze gene expression levels across all cells^[17]^, identifying diploid cells as normal cells, with normal and tumor cells separated due to differences in gene expression distributions. Subsequently, we classified tumor cells based on the previously obtained multi-omics consensus clustering approach, and used the “irGSEA” R package to perform pathway enrichment analysis on different subtypes, conducted metabolic differential analysis with the “scMetabolism” R package, and ultimately explored the interaction patterns of different subtype malignant cells with other cell populations using the “CellChat” package. In addition, we analyzed the scRNA-seq dataset GSE202642 using the same workflow as GSE229772, thereby validating important outcomes such as the robustness of the multi-omics clustering consensus typing method at the scRNA data level, the metabolic differences between different subtypes of malignant cells, and the variations in cell communication patterns.

## Results

### Construct multi-omics consensus clustering classification and perform validation

We utilized clustering prediction indicators, gap statistical analysis, silhouette scores, and insights from previous research involving 10 multi-omics integrated clustering algorithms to ascertain the number of subtypes (Fig.1A). Using a consensus integration method, we linked the clustering results to distinct molecular expression patterns of the transcriptome (mRNA, lncRNA, and miRNA), epigenetic methylation, and somatic mutations, thereby identifying two subtypes (CS1 and CS2) independently (Fig.1B). Significant differences in mRNA, lncRNA, miRNA, DNA CpG methylation sites, and mutation gene characteristics were observed between the two subtypes (Fig.1C). The five types of omics data used in the construction of the multi-omics consensus clustering classification can be found in the supplementary materials. This classification method is strongly correlated with overall survival (p < 0.001), with patients of the CS2 subtype exhibiting better prognostic survival compared to those of the CS1 subtype (Fig.1D). These characteristics suggest that our classification method can effectively distinguish between patients. Subsequently, employing the NTP method, we selected 200 genes specifically upregulated in each subtype as classifiers to validate the classification in an independent external validation set, ICGC, and found a high degree of consistency in the classification results. These 200 feature genes, along with the gene expression profiles and clinical data from the ICGC cohort that we used, can be found in the supplementary materials. Patients with CS1 and CS2 subtypes in the ICGC cohort exhibited similar prognostic differences (Fig.1E-G). This demonstrates the versatility and practical value of our classification method. We will explore the reasons for the significant prognostic differences between the two subtypes of patients by integrating multi-omics data. It will primarily focus on the sensitivity of tumors to current mainstream treatment options, including immunotherapy, targeted therapy, and chemotherapy.

**FIGURE 1.**
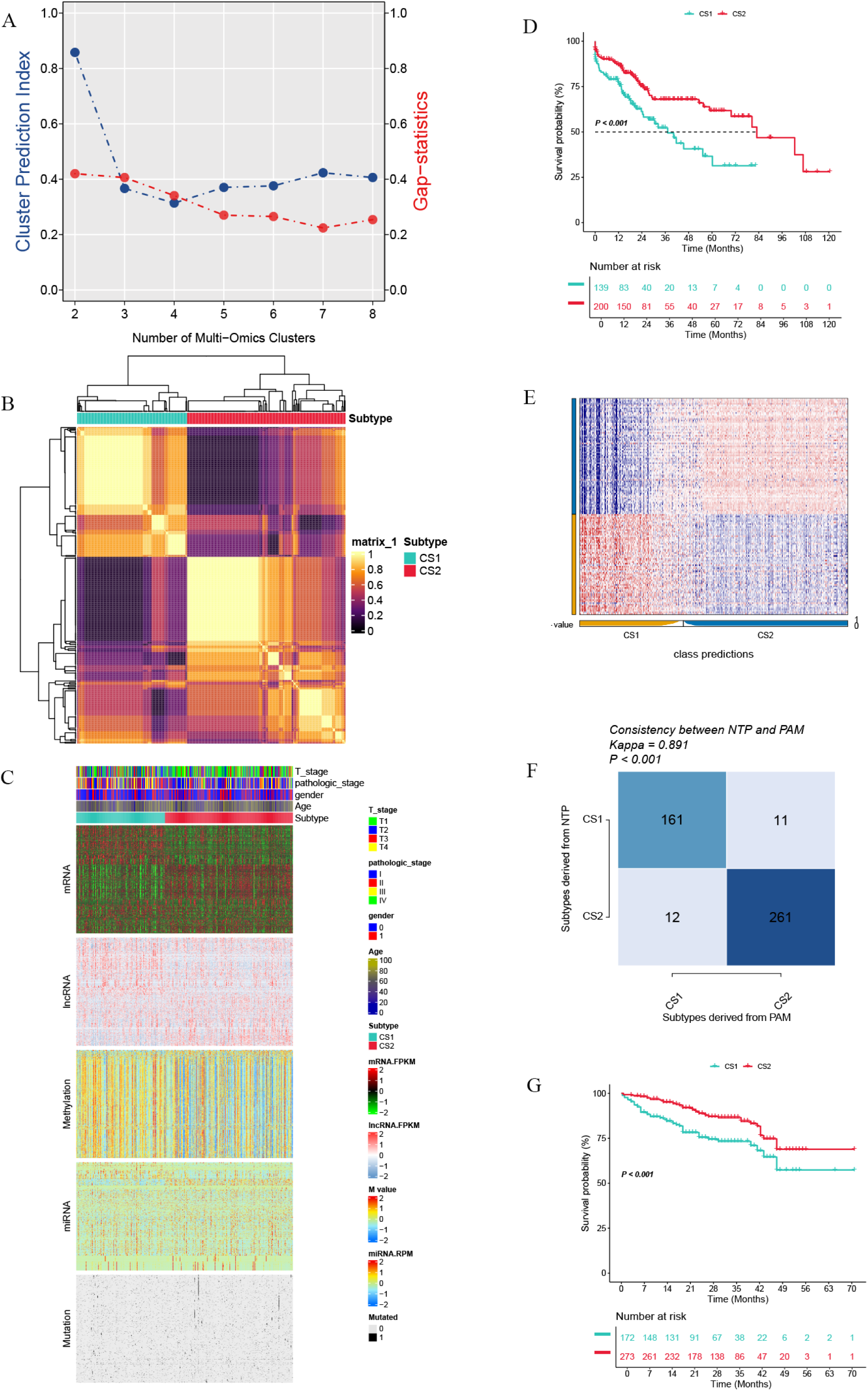
Multi-omics consensus clustering classification. (A) Calculation of optimal number of subtypes based on cluster prediction index and gap-statistics. (B) Heatmap of the consistency of the integration of the results of the 10 typing algorithms. (C-D) Significant multiomics characterization and prognostic differences between subtypes. (E-G) Validation of consistency of typing results and prognostic value in an external cohort.

### Molecular Characteristics of Multi-Omics Consensus Clustering Subtypes

Based on the correlation between tumor mutation burden (TMB) and immune therapy response, we analyzed the gene mutation landscape of two subtypes and found that both subtypes exhibited a high frequency of mutations in TP53, KMT2D, and DNAH6 genes(Fig.2A). Then we analyzed the TMB of the two subtypes of patients to assess the impact of immune therapy responsiveness on the prognosis of these subtypes. The results showed no significant difference in TMB between the two patient subtypes. (Fig.2B-C). Subsequently, we analyzed the bulk RNA sequencing data of patients within the two subtypes, identified differentially expressed genes, and conducted enrichment analysis on these genes. The results revealed significant differential enrichment in several biological pathways, including those related to metabolism and signal transduction, such as lipid peroxidation, fatty acid metabolism, and amino acid metabolism, suggesting notable biological differences between the tumors and associated tissues of the two subtypes (Fig.2D-E). The list of differentially expressed genes can be found in the supplementary materials. Previous studies have reported that heterogeneity among different tumor tissues of the same tumor type largely stems from metabolic reprogramming and changes in signal transduction pathway activity^[18–19]^. This implies that the heterogeneity between the tumor tissues of the two subtypes may also arise from these factors. Finally, we compared the consistency of subtyping methods with the pathologic staging of patients(Fig.2F).

**FIGURE. 2.**
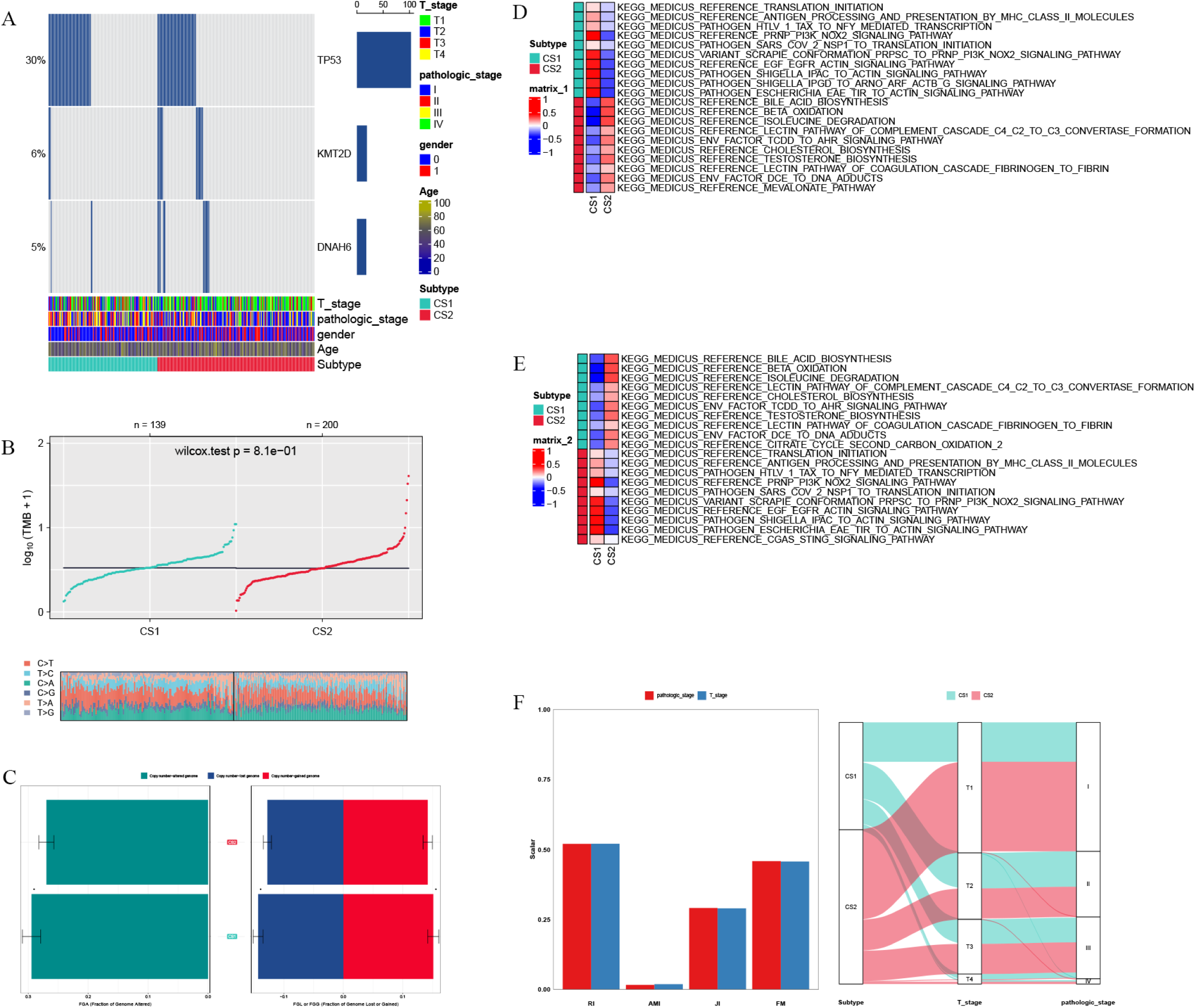
Differences in molecular characterizaiion of subtypes. (A) Differences in mutation landscapes of subtypes. (B-C) Genomic characterization from TMB and FGA of subtypes. (D-E) Functional emichment of up and down-regulated differential genes between subtypes. (F) Concordance between subtypes and patients’ pathologic staging.

### Cell, signaling pathways, stemness, and other phenotypic characteristics of multi-omics consensus clustering subtypes

In the results presented in Fig. 2, we found that there may be no significant difference in tumor mutation burden between the two subtypes. Therefore, we analyzed the tumor immune microenvironment factors that have been frequently reported to potentially affect the response of tumors to immunotherapy. We applied the ssGSEA algorithm to assess the enrichment status of immune-related cells in the tumor microenvironment (TME) of patients from the TCGA cohort, based on a gene set of common immune-related cell characteristics compiled from multiple literature sources. (Fig.3A). The gene set utilized for this analysis was compiled and summarized from various literature sources. After the preliminary analysis of the immune response of the two tumor subtypes to immunotherapy, we utilized the drug intervention data from the pRRophetic database to assess the sensitivity of the two tumor subtypes to common liver cancer therapies, including chemotherapy and targeted therapy. We found the CS2 subtype exhibited higher sensitivity to 5-fluorouracil, Gefitinib, Afatinib, and Erlotinib compared to the CS1 subtype (Fig.3B). Among these, 5-fluorouracil is the main drug used in hepatic arterial infusion chemotherapy for hepatocellular carcinoma (HCC) patients, whereas Gefitinib, Afatinib, and Erlotinib are tyrosine kinase inhibitors (TKIs) that primarily inhibit epidermal growth factor receptor (EGFR). We further compared the level of infiltration of each cell taxon in subtypes in the TME(Fig.3C). Within the TME of the CS2 subtype, associated with a comparatively more favorable prognosis, nearly all assessed cell distribution levels demonstrated a decrease in comparison to those in the CS1 subtype. This encompasses traditionally recognized inhibitory immune cells like MDSCs and Regulatory T cells, alongside pivotal tumor-killing immune cells like Activated CD8+ T cells and NK cells. In addition, we also conducted similar immune infiltration assessments using ESTIMATE and EPIC scores. The results indicated that the findings from these methods were consistent with those obtained using ssGSEA (Fig. S1A-B). These findings may imply reduced levels of tumor-associated antigens within CS2 tumor tissue, leading to a comparatively subdued activation and induction of immune cells within the TME. After discovering significant differences in the responses of the two tumor subtypes to mainstream liver cancer treatment methods, we employed the ssGSEA method, combined with gene sets corresponding to pathways, to preliminarily assess several common pathways that may influence the tumor’s response to treatment. We discovered that the CS1 subtype is more active than the CS2 subtype in multiple pathways associated with tumor development and progression, including the MAPK, JAK-STAT, NFκB, PI3K-AKT, and VEGF signaling pathways, as well as pathways involved in angiogenesis and hypoxia. No significant differences were observed in the cAMP signaling pathway or the mRNAsi stemness score (Fig. 3D).

**FIGURE 3.**
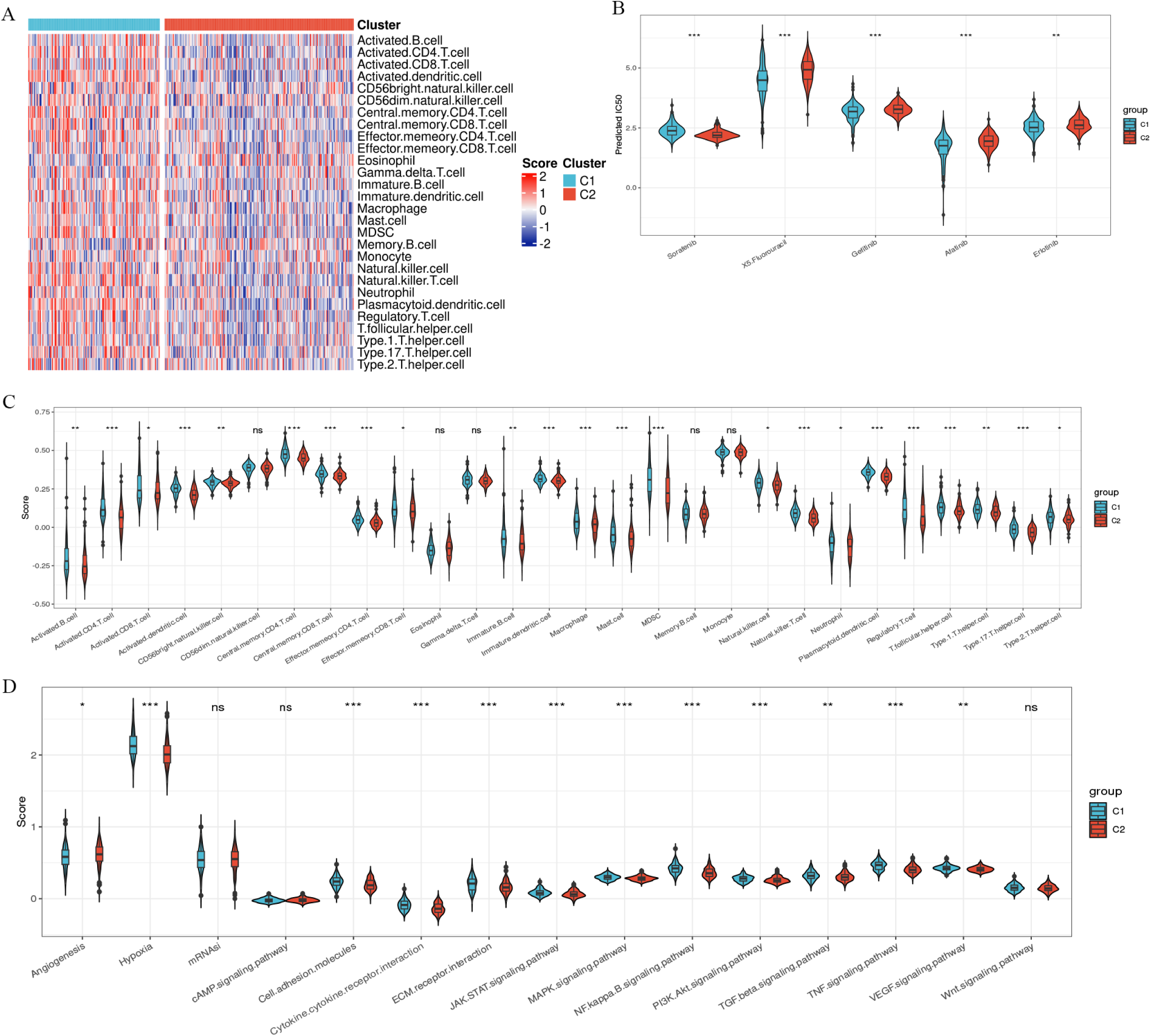
Phenotypic featmes of subtypes that may be associated with tumor development. (A) Landscape of subtype Th1E immune infiltration. (B) Differences in subtype responsiveness to different drugs. (C) Comparison of immune cell infiltration levels in subtypes ofTME. (D) Comparison ofhepatocellular carcinoma-related signaling pathways and biological process activity in subtypes.

### Constructing a prognostic prediction model based on the consensus clustering of multi-omics features

After obtaining a classification method with clinical prognostic predictive value and gaining some theoretical support from RNA bulk data analysis, we constructed a patient prognosis prediction model based on this classification method to achieve a more accurate and reasonable prediction of patient outcomes. We selected the top 30 most significantly differentially expressed genes from CS1/CS2 gene templates as candidate prognostic risk genes. Subsequently, we applied 101 algorithmic approaches to determine the correlation and impact weights between these 30 genes and prognosis, leading to the construction of a prognostic prediction model. Simultaneously, we calculated the average C index of each model to assess the predictive capability across all models. The numbers represented by the blue bars indicate the average C-index of the model in both the training and internal validation sets (Fig. 4A). Subsequently, we obtained the risk coefficients of the five genes used to build the RSF model by stepwise Cox regression analysis (Fig.4B). We then employing this model to evaluate patients’ prognostic risk within TCGA-LIHC cohort and stratifying them based on the optimal cutoff value, we observed a notably poorer prognosis among high-risk group patients compared to their low-risk counterparts. The 5 central genes, and the Hazard Ratio associated with these genes in the TCGA-LIHC cohort is depicted in Fig. 4C. Additionally, we illustrated the patients’ prognostic outcomes and variations in central genes as the prognostic risk escalated (Fig. 4D). Subsequently, we validated the model using independent external validation set, the ICGC and GSE14520 cohorts. A notable disparity in prognosis between the two groups was evident, aligning with the outcomes observed in the TCGA-LIHC cohort (Fig. 4E-F). In addition, we also plotted 1-, 2-, 3-year AUC curves in the TCGA-LIHC, ICGC and GSE14520 cohorts (Fig. S1C-E).

**FIGURE 4.**
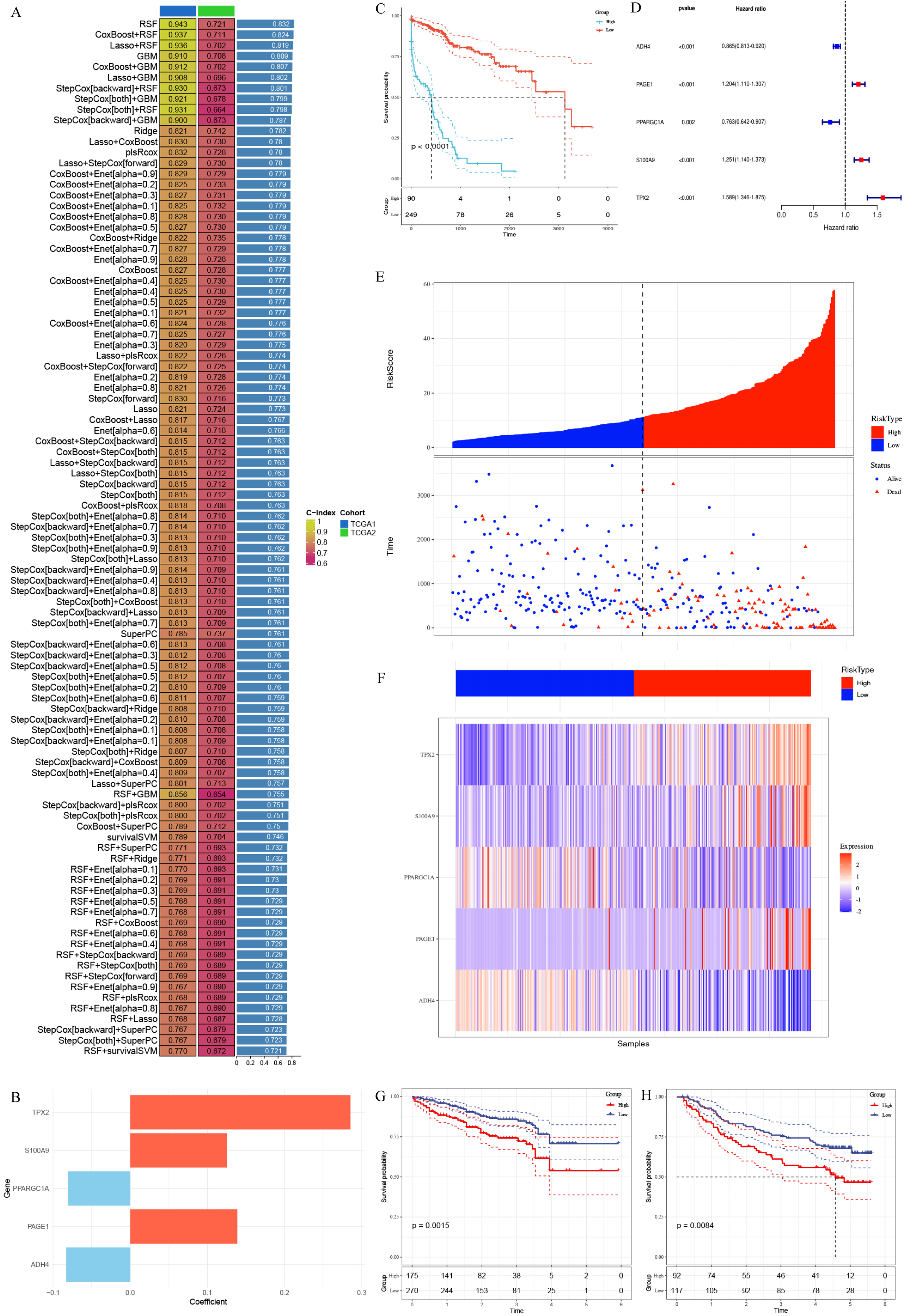
Building prognostic prediction models through machine learning. (A) 101 machine learning algorithms to build screening optimal prognostic models(TCGA1=training set,TCGA2=internal validation set). (B) Regression coefficients of 5 genes obtained in stepwise Cox regression.

### Comparison of Prognostic Prediction Models

With the rapid advancement and widespread adoption of Next Generation Sequencing (NGS) technology, numerous prognostic prediction models based on gene expression have been extensively reported. After developing our model, we reviewed and collected prognostic prediction models reported in the past five years and included 19 of these models, which are based on various foundations such as immune infiltration, glucose metabolism, and lipid metabolism. We calculated the concordance index for a total of 20 models in both TCGA-LIHC dataset and ICGC dataset. The average C-index of these models was derived to evaluate the prognostic predictive performance of the model. The results indicated that our model demonstrated superior prognostic prediction performance compared to all other models in both cohorts (Fig. 5A). Subsequently, we incorporated clinical features related to prognosis into the model to construct a nomogram (Fig. 5B). The calibration curve demonstrated that the nomogram had predictive accuracy more consistent with actual conditions (Fig. 5C). Decision curve analysis (DCA) indicated that both the nomogram and the original model could benefit patients, with the nomogram performing slightly better (Fig. 5D).

**FIGURE 5.**
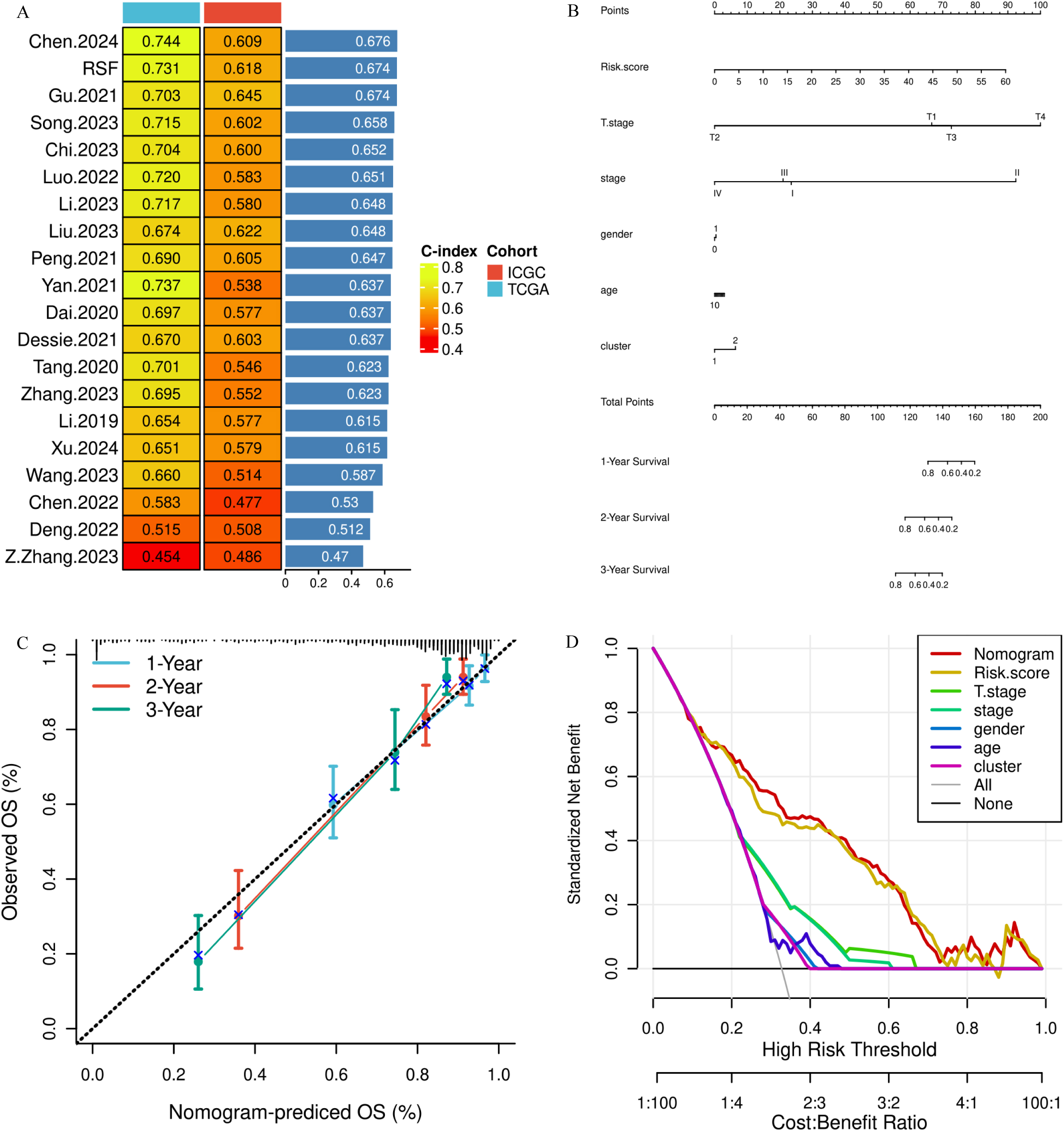
Comparison and refinement of prognostic prediction models. (A) Comparison of predictive performance of prognostic models with some prognostic models published in the last 5 years. (B) Constructing a nomogram incorporating patient clinical characteristics. (C-D) Evaluating model performance based on calibration curve and decision curve analysis.

### scRNA-seq Data Processing

Given the significant prognostic differences between patients with CS1 and CS2, as well as the notable heterogeneity observed at the cellular and molecular levels in tumor tissues from the RNA-Bulk data analysis, we attempted to further explore the potential mechanisms underlying these differences between the two subtypes by analyzing scRNA-seq data. The GSE229772 dataset from the GEO database was chosen for our analysis. Following quality control filtering, 56,937 cells were derived from 31 samples. By employing hierarchical and unsupervised clustering methods, we delineated cell subclusters according to their expression profiles, categorizing the cells into 13 subclusters (Fig. 6A). Subsequently, we employed established marker genes to profile various cell types and assign annotations, discerning 12 primary cell subgroups: B cells, stromal cells, vascular endothelial cells, CD8+ T cells, epithelial cells, fibroblasts, germ cell-like cells, hepatic stellate cells, hepatocytes, macrophages, neutrophils, and T cells (Fig. 6B-D). Within these, the immune cell group prevailed as the most abundant, succeeded by stromal cells and fibroblasts (Fig. 6E).

**FIGURE 6.**
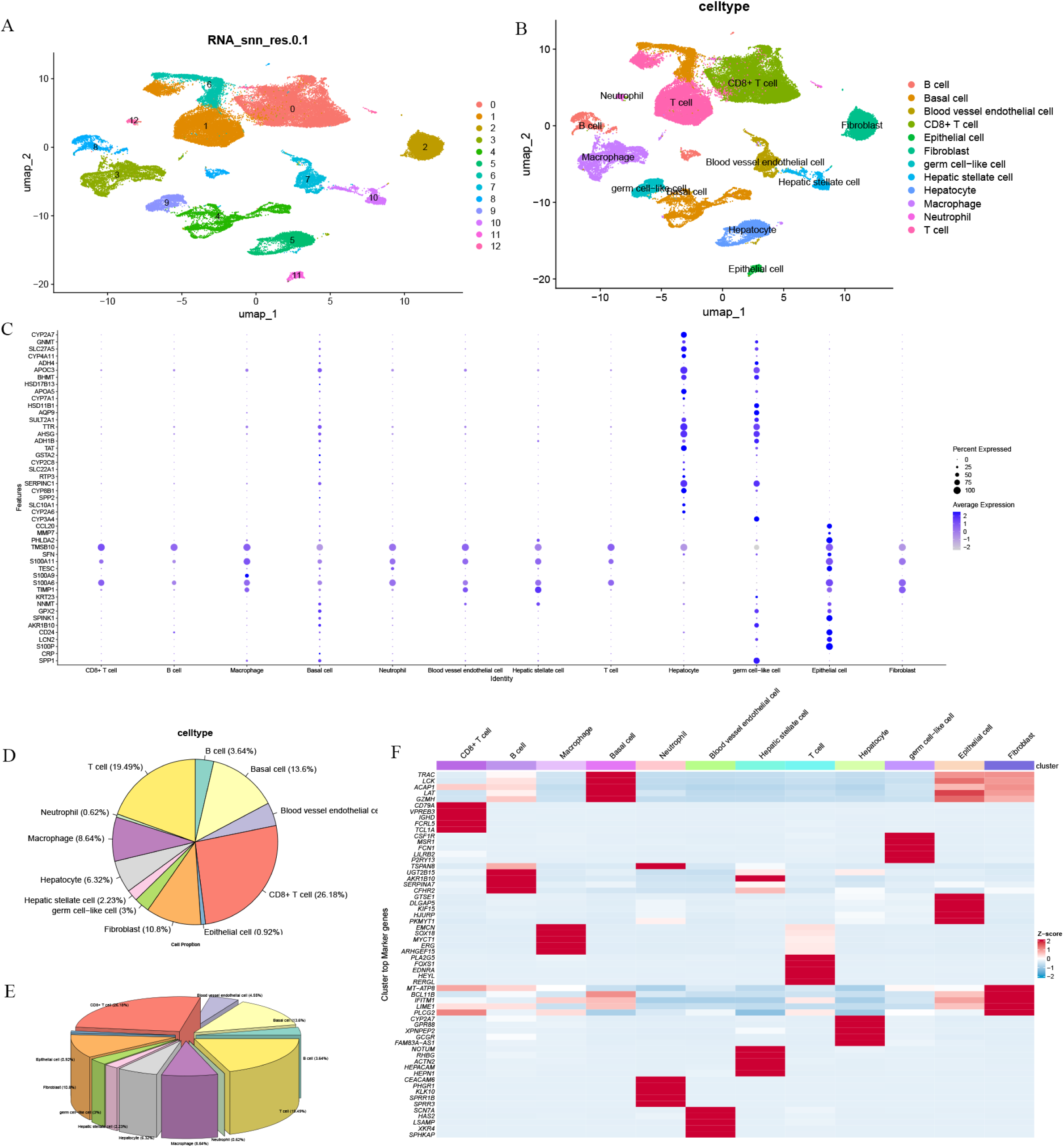
ScRNA-seq cell s01iing and annotation. (A-C) Cellular subpopulation delineation and annotation based on cellular characterization genes. (D-E) Visualization of the proportions of each cell subgroup. (F) Differentially expressed genes by cellular subpopulations.

### Characteristic Analysis of Different Cell Subpopulations

To investigate pathway heterogeneity among different cell subclusters, we identified differentially expressed characteristic genes within these subclusters. Subsequently, we employed multiple algorithms, including AUCell, UCell, singscore, ssgsea, JASMINE, and viper, to assess pathway activity variation. Differential analysis results were comprehensively evaluated using robust rank aggregation (RRA), and gene sets significantly enriched across multiple enrichment analysis methods were filtered to obtain consensus results. Statistically significant differentially enriched pathways from these consensus results were visualized (Fig.7A-B). We observed significant differences in several metabolic and signaling pathways among different cell subclusters. Given the reported impact of metabolic changes and signaling pathway variations on the TME and their implications for heterogeneity, we specifically analyzed these pathways. Utilizing the VISION algorithm, we assessed metabolic levels across each cell subcluster and visualized the differences in a dot plot (Fig.7C). Our findings indicated pronounced metabolic variations among cell subclusters, with germ cell-like cells and hepatocytes notably exhibiting heightened metabolic activity compared to other cell groups. Subsequently, we utilized the CellChat R package’s integrated databases for secreted signaling, ECM receptor signaling, cell-cell contact, and non-protein signaling to explore cell interactions within the single-cell dataset. Initially, we examined the cell communication landscape between distinct cell groups and constructed a cell communication network (Fig.7D). Our analysis revealed robust communication links between various cell groups, particularly notable between fibroblasts and hepatic stellate cells, which exhibited signaling connections with nearly all other cells. Then, we visualized the signaling output and input patterns (Fig.7E-G) to illustrate the contributions of different cell groups at both ends.

**FIGURE 7.**
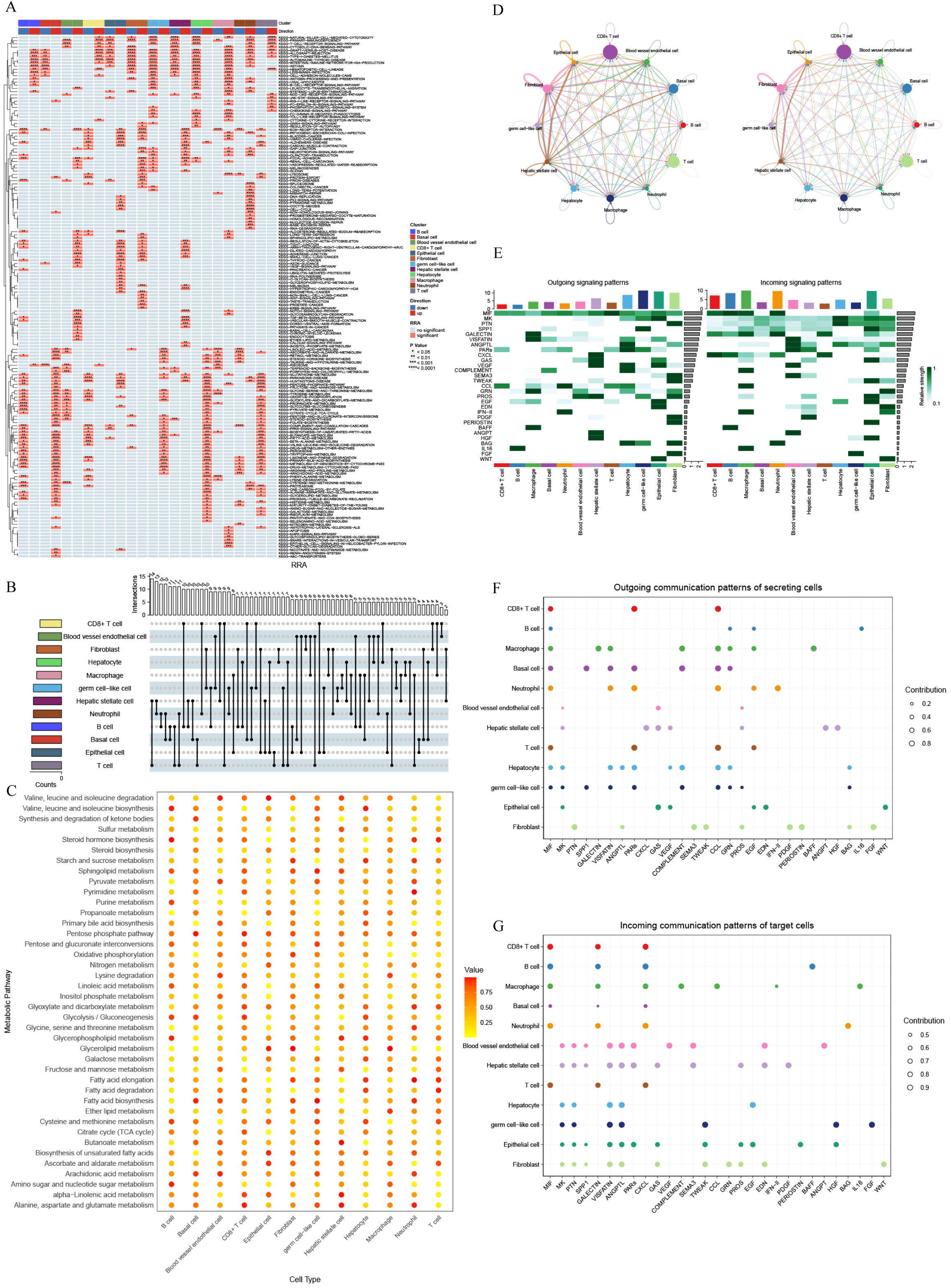
Phenotypic characterization of cellular subpopulation. (A-B) Functional enrichrnent analysis of differentially expressed genes within cell subpopulation. (C) Analysis of the metabolic landscape of cell subpopulations. (D) Cellular polymeric communication network. (E) The contribution of signal strength in both incoming and outgoing signals shapes the landscape of cellular subgroups. (F-G) Analysis of signal transmission into and out of cellular subgroups.

### Characteristics of Different Subtypes of Malignant Cells at the Single-Cell Level

To delve deeper into the heterogeneity of the two tumor subtypes, we initially utilized the copykat algorithm to pinpoint potential malignant cell subclusters, including hepatocytes, fibroblasts, and hepatic stellate cells, by detecting whole-genome aneuploidy. This approach facilitated the identification of potential malignant cells, which were subsequently aligned with the prior cell annotation results (Fig.8A). Subsequently, we employed the previously derived multi-omics consensus clustering features to classify the malignant cells into two subtypes, CS1 and CS2, by NTP method (Fig.S2). This analysis enabled the identification of differentially expressed genes between the two subtypes. Enrichment analysis of these genes revealed significant differences in various signaling pathways between CS1 and CS2 malignant cells, including the PI3K-AKT and P53 signaling pathway (Fig.8B). A targeted analysis of the metabolic variances between the CS1 and CS2 subtypes revealed notable differences, indicating significantly higher metabolic activity in CS1 compared to CS2 (Fig.8C). Then, upon integrating the malignant cells classified as CS1 and CS2 into the comprehensive cell communication network (Fig.S3), we observed that CS1 and CS2 exhibited widespread regulation of all other cell groups in the TME (Fig.8D). Subsequently, we analyzed the signal output and input patterns of various cellular subgroups and found that the MIF signaling pathway exhibited the highest frequency and greatest contribution in the signal communication network, whether at the input or output end(Fig.8E-G). We further analyzed the output levels of the MIF signaling pathway in various cellular subgroups, focusing on which cells were outputting the MIF signal. The results showed that the CS1 and CS2 subtypes mainly output MIF, with CS1 subtype being relatively predominant(Fig.8H). Further investigation into the regulatory effects of CS1 malignant cells on other cell groups demonstrated that the CS1 subtype exerted extensive regulatory effects on nearly all immune cells in the TME, including CD8+ T cells, macrophages, B cells, and neutrophils, primarily through the MIF- (CD74+CXCR4) pathway (Fig.9). It is noteworthy that during our analysis of GSE202642, we found that the metabolic differences and cell communication pattern variations among different subtypes of malignant cells were largely consistent with the results obtained from the analysis of GSE229772 (Fig.S4-6).

**FIGURE 8.**
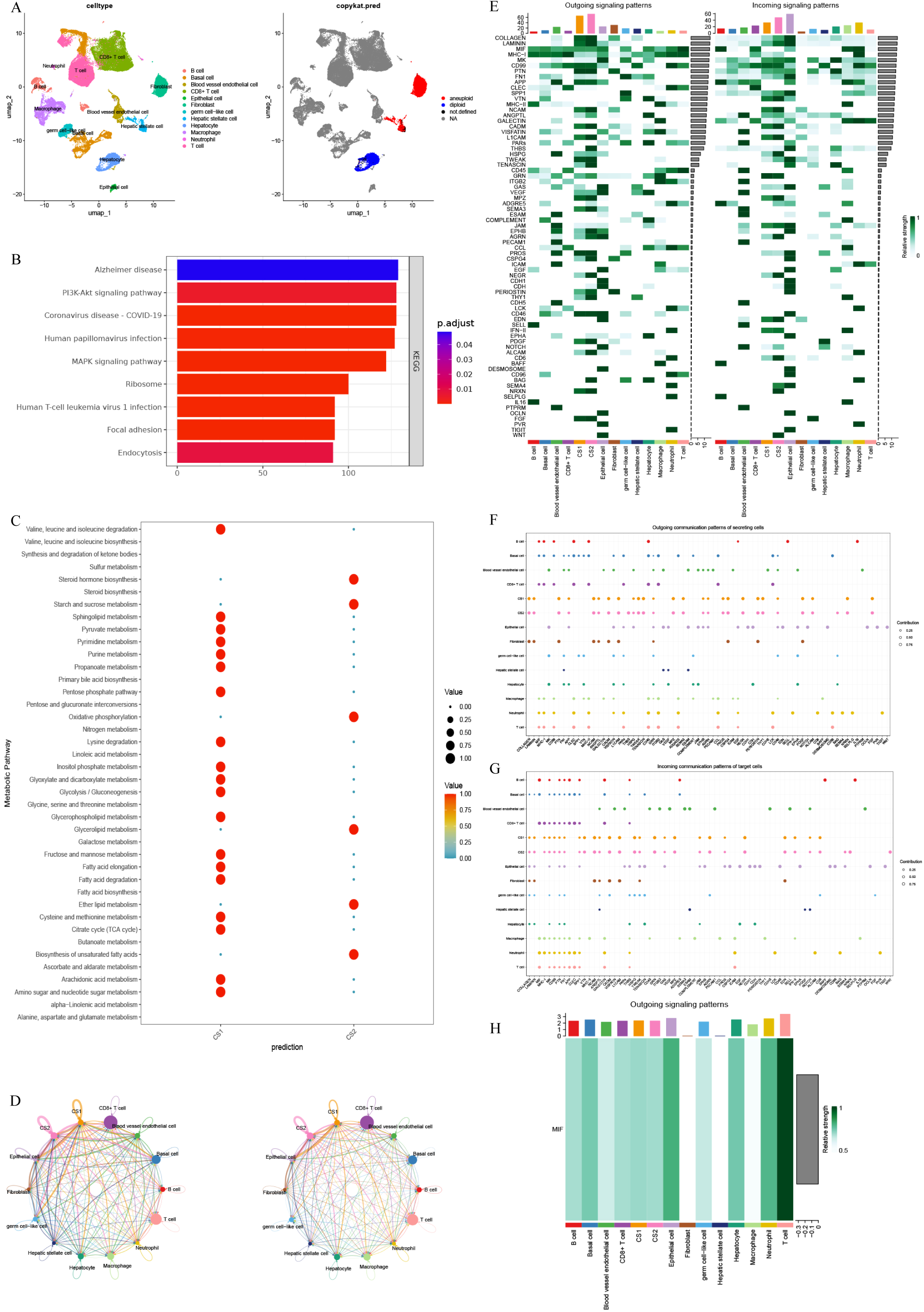
Exploring the characteristics of two subtypes of malignant cells at the scRNA-seq level. (A) Mapping of malignant cells in the cell clustering results. (B) Enrichment analysis of differential genes between different subtypes of malignant cells. (C) Metabolic level analysis of different subtypes of malignant cells. (D) Annotation of the cellular subpopulation signaling landscape after malignant cell annotation. (E-G) Analysis of signal input-output patterns in cellular subpopulation. (H) Differential levels of the MIF signaling pathway in cellular subpopulation outputs.

**FIGURE 9.**
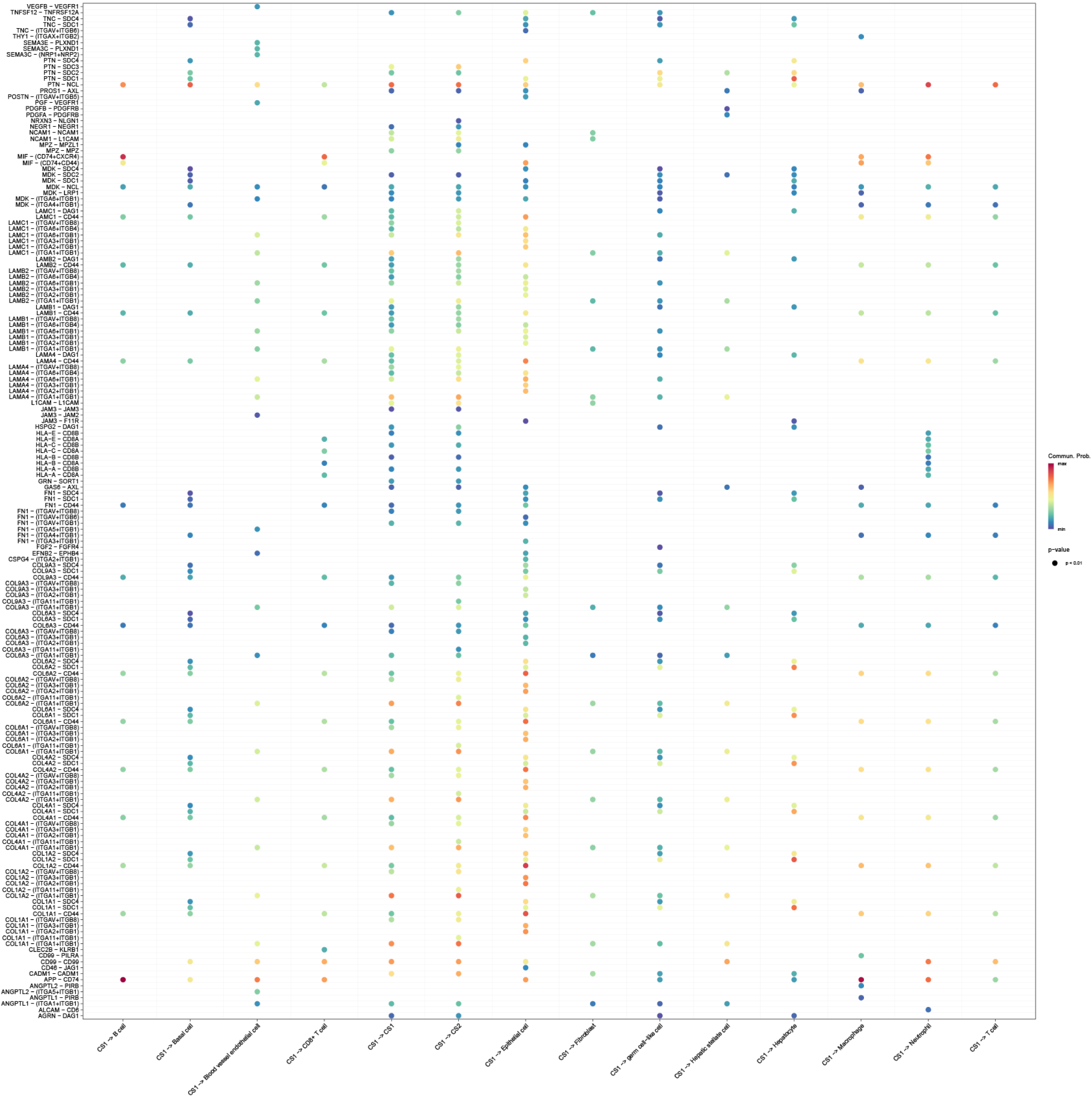
Signaling modulation pattern of CSl subtype malignant cells on other cells.

## Discussion

The role of genes in disease development, particularly in tumorigenesis, is well-established. Studying gene expression processes significantly enhances our understanding of tumors. Gene expression is intricately regulated by genetic mechanisms such as methylation and mutations. Therefore, integrating multi-omics data analysis is crucial for gaining deeper insights into disease-specific regulatory mechanisms. However, most studies to date have primarily focused on single omics, such as mRNA and miRNA expression research. Moreover, the choice of omics clustering methods in research often relies on personal preference, exacerbating the limitations of individual approaches in studies. Our research aims to bridge this gap by integrating ten state-of-the-art clustering algorithms to identify two prognostic subtypes with distinct characteristics, which could potentially facilitate precise stratification and treatment of HCC patients. Furthermore, our novel classification method has demonstrated stability across multiple cohorts.

Significant prognostic differences among patients with the same tumor type may be influenced by various factors, including tumor tissue heterogeneity, metabolic reprogramming, signaling patterns, and disparities in the TME. As such, our study undertook initial investigations to elucidate the underpinnings of these prognostic discrepancies across patient subtypes. Initially, we evaluated the mutation landscape across subtypes and observed no notable differences in TMB and FGA between the two subtypes, implying that subtype heterogeneity may not originate from gene mutations. Subsequent analysis of differentially expressed genes and gene enrichment indicated significant enrichment in tumor-related pathways, particularly metabolism and signal transduction pathways like lipid peroxidation, fatty acid metabolism, amino acid metabolism, PI3K-NOX, and EGF-EGFR signaling pathways. This suggests potential variances in metabolic reprogramming and signaling patterns between the subtypes, which can influence patient prognosis by governing the TME. Consequently, we conducted a further analysis of immune cell infiltration in the TME, revealing lower immune cell infiltration in CS2 compared to CS1, implying a potentially lower malignancy in CS2 with reduced release of tumor-associated antigens and subsequent lower immune response and infiltration. Additionally, we thoroughly examined the signaling pathways and biological characteristics closely associated with tumor development. The findings revealed that the CS1 subtype exhibited greater activity than the CS2 subtype in multiple pathways linked to tumorigenesis, including the MAPK, JAK-STAT, NFκB, PI3K-AKT, VEGF, angiogenesis, and hypoxia pathways. These pathways are widely recognized for their close association with tumor development and their significant roles in tumor transformation and progression. Furthermore, the hypoxia pathway and angiogenesis pathway are well-documented to contribute to the formation of an immunosuppressive TME in HCC^[20]^. Hypoxia directly induces VEGF production^[21–22]^, which can exert inhibitory effects on various immune cells, such as CD8+ T cells^[23]^. Additionally, VEGF-induced angiogenesis within the TME exacerbates hypoxia, leading to a detrimental feedback loop^[24]^. These factors likely contribute to the observed heterogeneity and prognostic differences between the two subtypes. Finally, we conducted an in-depth assessment of the impact of subtyping on clinical treatment and prognosis assessment for patients. Our findings revealed that CS2 exhibited enhanced sensitivity to the conventional drug 5-fluorouracil utilized in hepatic arterial infusion chemotherapy, as well as to several tyrosine kinase inhibitors, including gefitinib, afatinib, and erlotinib, compared to CS1. This discovery holds the potential to inform tailored treatment strategies for patients based on their subtype, consequently improving treatment efficacy and prognosis.

The above findings suggest that our classification method reliably distinguishes patients into two subtypes characterized by significant prognostic differences. These prognostic disparities may arise from heterogeneity linked to variations in metabolic reprogramming and signaling patterns, thus providing a theoretical foundation for the classification’s validity. Consequently, we developed a prognostic prediction model based on subtype characteristics to enhance the clinical applicability of the classification and effectively predict patient prognosis. Machine learning algorithms serve as powerful tools for analyzing multi-omics data and constructing prognostic models. We utilized the TCGA-LIHC cohort as the training and validation set. After testing 101 algorithm combinations to identify the optimal model, we employed the C-index to evaluate predictive performance, ultimately determining the model with the highest C-index, which signifies superior predictive capability. In addition to selecting the best results from 101 algorithm combinations to achieve greater accuracy and reliability, the use of the LOOCV method can significantly reduce the bias and randomness that arise during the application of the training and validation sets, compared to previous methods that directly split the training and validation sets by a certain ratio. The data revealed that the prognostic model, developed using the Random Survival Forest (RSF) algorithm and incorporating 5 hub genes, demonstrated superior performance. Upon assigning risk scores and stratifying patients in both the training and validation sets based on this model, it became evident that individuals in the high-risk category exhibited significantly poorer prognoses compared to those in the low-risk category, affirming the success of our model construction. Subsequently, we compared our model with 19 previously established prognostic prediction models to predict patient outcomes and assessed their performance in both cohorts. Through comprehensive evaluations, our model consistently outperformed the other models across both sets. Additionally, we conducted further assessments using calibration curves and decision curve analysis, yielding consistent results indicating the robust predictive capabilities of our model for patient prognosis. It should be noted that although the model building method we used has obvious advantages compared with some early modeling methods, it still belongs to the mathematical analysis based on public databases. Therefore, during the analysis process, the mathematical logic of the algorithm inevitably obscures some factors that may be clinically relevant to the prognosis of patients.

After exploring the clinical implications of our classification method, we conducted an analysis of tumor subtype heterogeneity at the single-cell level to uncover potential mechanisms contributing to differences in prognosis and heterogeneity between subtypes. Initially, we annotated a single-cell sequencing dataset comprising tumor tissue samples from 31 HCC patients, effectively categorizing cells into 12 distinct groups. Subsequently, leveraging the CopyKAT algorithm, we analyzed gene expression levels to identify non-diploid cells as malignant. Employing the same classification method used previously, we successfully categorized malignant cells into two subtypes and assessed differential gene expression and metabolic pathways between them. Our findings aligned with RNA-bulk sequencing data, highlighting significant distinctions between subtypes across various tumor-related signaling pathways and metabolic processes.

Our findings indicate significant heterogeneity among tumor cells of the two subtypes. Additionally, we conducted an analysis of signaling patterns between tumor cells and other cell groups. The results revealed that CS1 malignant cells exert substantial regulatory effects on immune cells within the TME, including CD8+ T cells, macrophages, B cells, and neutrophils, mediated by the MIF-(CD74+CXCR4) pathway. Prior research has elucidated that MIF binds to its receptor CD74, leading to its phosphorylation and the recruitment of CD44, triggering downstream phosphorylation signals via the proto-oncogene SRC^[25–26]^. This cascade encompasses the transcriptional activation of cyclin D1, phosphorylation of retinoblastoma protein, and the involvement of multiple pro-proliferation and anti-apoptotic pathways such as the MAPK/ERK and PI3K/AKT pathways^[27–28]^.

The MIF signaling pathway is intricately involved in diverse cellular processes. It stabilizes the p53-MDM2 complex, promoting cell proliferation and inhibiting apoptosis^[29]^. There is a synergistic relationship between MIF and HIF-1α, where HIF-1α binds to the 5’-UTR hypoxia response element of MIF, thereby enhancing MIF expression regulated by CREB^[30]^. Additionally, MIF activates a p53-dependent feedback loop of HIF-1α, crucial for tumor hypoxia adaptation^[31]^. Moreover, MIF prevents proteasome-mediated degradation of HIF-1α and forms a ternary complex with CSN5 and HIF-1α, amplifying the tumor hypoxia response^[32]^. Our prior findings indicate heightened activity in MAPK, PI3K-AKT signaling, and the hypoxia pathway in the CS1 subtype compared to CS2. Integrating RNA-bulk and single-cell sequencing data suggests a pivotal role for the MIF pathway in subtype heterogeneity.

We have reviewed and summarized several articles that have been published regarding the classification of HCC patients and the construction of prognostic models using machine learning. A considerable number of studies focus on specific pathways that may affect the tumor microenvironment, metabolic reprogramming, or patient prognosis (such as fatty acid metabolism, ferroptosis, and autophagy), using the expression levels of genes in these pathways as classification features for HCC patients and constructing prognostic models based on general regression analysis. Although this research method can produce classification results related to patient prognosis and generate prognostic models, it also has some issues. First, focusing on a specific pathway from the outset of the study may exclude most classification features outside the designated pathway, leading to limitations in the research. Second, these studies only use the mRNA expression levels of genes to construct classification features, making the utilization of omics data rather singular, which may affect the robustness of the classification method. However, a positive aspect is that some studies have selected key genes associated with prognosis and conducted experimental validation, which is helpful for exploring how classification methods can effectively distinguish patient prognosis.

In comparison to prior studies, our research exhibits several distinctive characteristics. First, we acknowledged the potential substantial heterogeneity in HCC patients, shaping our research perspective towards offering more precise stratification and treatment for patients. Second, we incorporated 5 types of omics data and employed 10 clustering methods for analysis, ultimately amalgamating them to derive a consensus clustering method, thereby mitigating the impact of clustering method selection bias on our analysis. Third, our modeling was underpinned by the features of consensus clustering subtypes, utilizing 10 machine learning algorithms combined into 101 algorithmic configurations for modeling. We identified the model with the best average C-index performance, surpassing several previously reported prediction models in terms of predictive efficacy. Fourth, subsequent to identifying potential heterogeneity between subtypes in RNA-bulk data analysis, we further delved into single-cell data. Fifth, we employed various methods to validate the key results, including the validation of multi-omics profiling features using the ICGC cohort and two scRNA-seq datasets, as well as validating the key results obtained from the scRNA-seq data analysis with GSE202642, which will significantly strengthen the robustness of our results. Our exploration revealed the involvement of the MIF signaling pathway in tumor heterogeneity and the resulting prognostic disparities in HCC patients. This comprehensive approach allowed us to conduct an extensive study around the consensus clustering method, elucidating both its clinical application value and the mechanisms underpinning the observed heterogeneity and prognostic differences between subtypes.

## Conclusion

In summary, our study is based on a new multi-omics clustering consensus typing method. While discovering the prognostic value of this typing method through the analysis of RNA-bulk data and scRNA-seq data, we focused on the tumor heterogeneity caused by metabolic reprogramming and changes in signaling transduction patterns. We initially explored the reasons for the differences in prognosis among patients of different types and found that the MIF signaling pathway is involved in the occurrence and development of HCC. Due to the use of bioinformatics and limited samples and survival data in our study, it is necessary to further validate potential mechanisms through subsequent basic experiments and analysis of larger cohort data.

## Acknowledgments

This work was supported by the National Natural Science Foundation of China under grant 82272902, the Natural Science Foundation of Hubei Province of China under grant 2024AFD424 and the Beijing Public Health Foudation under grant 2023158.

## Data Availability Statement

The data analyzed in this study were obtained from The Cancer Genome Atlas(TCGA) at TCGA-LIHC, International Cancer Genome Consortium(ICGC) at LIHC-JP, Gene Expression Omnibus (GEO) at GSE229772 and GSE202642.

