## Supplementary Figure for "Decoding Liver Cancer Prognosis: From Multi-omics Subtypes, Prognostic Models to Single Cell Validation"

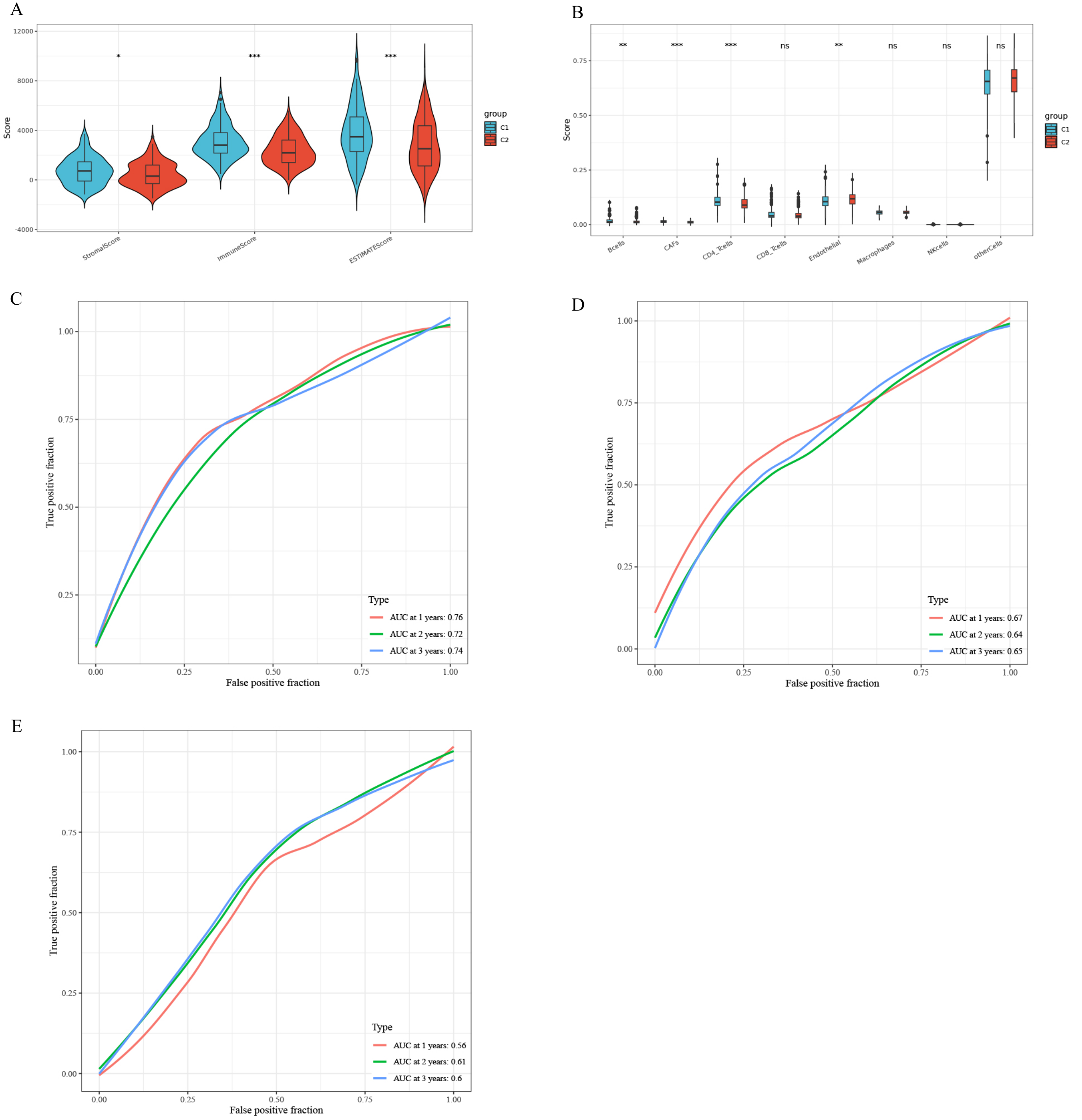


**Supplementary Figure 1**. (A) Comparison of immune landscape in subtypes of TME by ESTIMATE score. (B) Comparison of immune landscape in subtypes of TME by EPIC score. (C-E) The 1-, 2-, and 3-year AUC curves of TCGA-LIHC, ICGC, and GSE14520 cohorts.

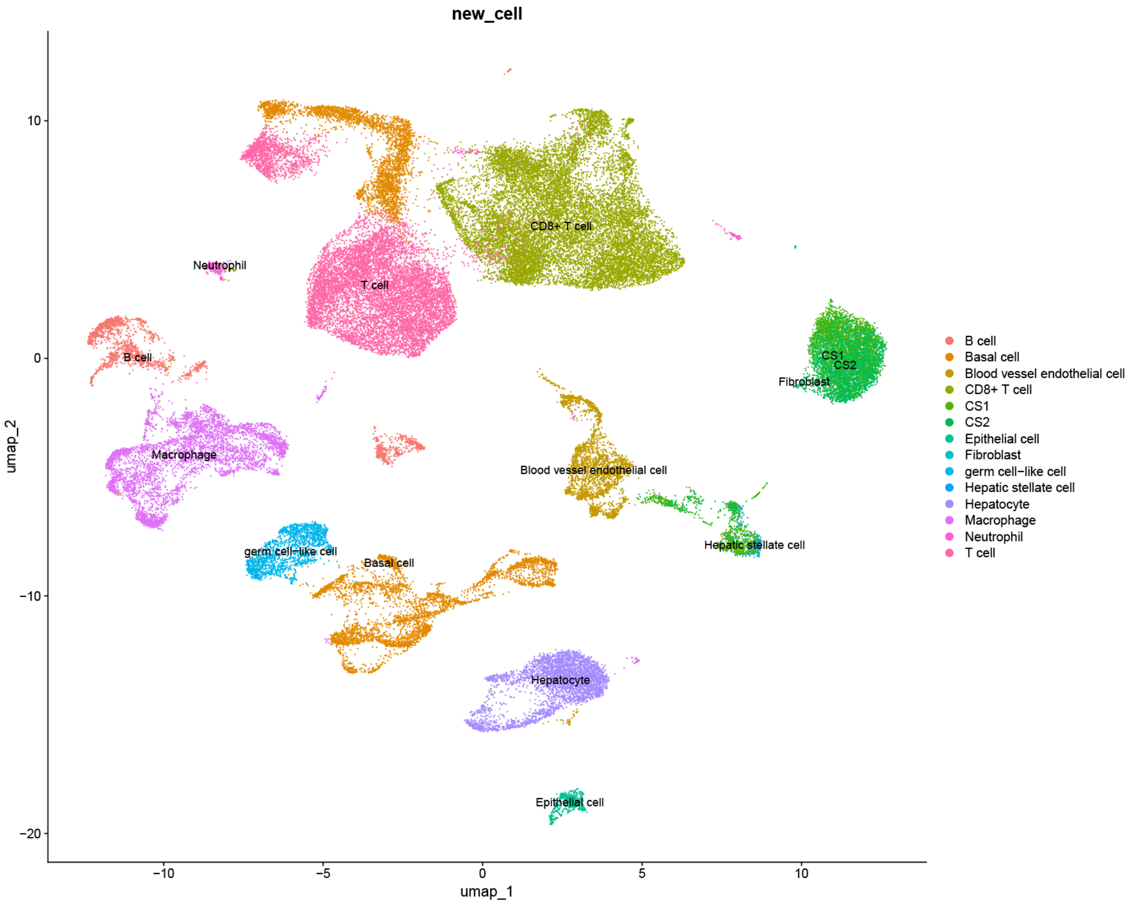


**Supplementary Figure 2.** The locations of the two subtypes of malignant cells in the overall cell population.

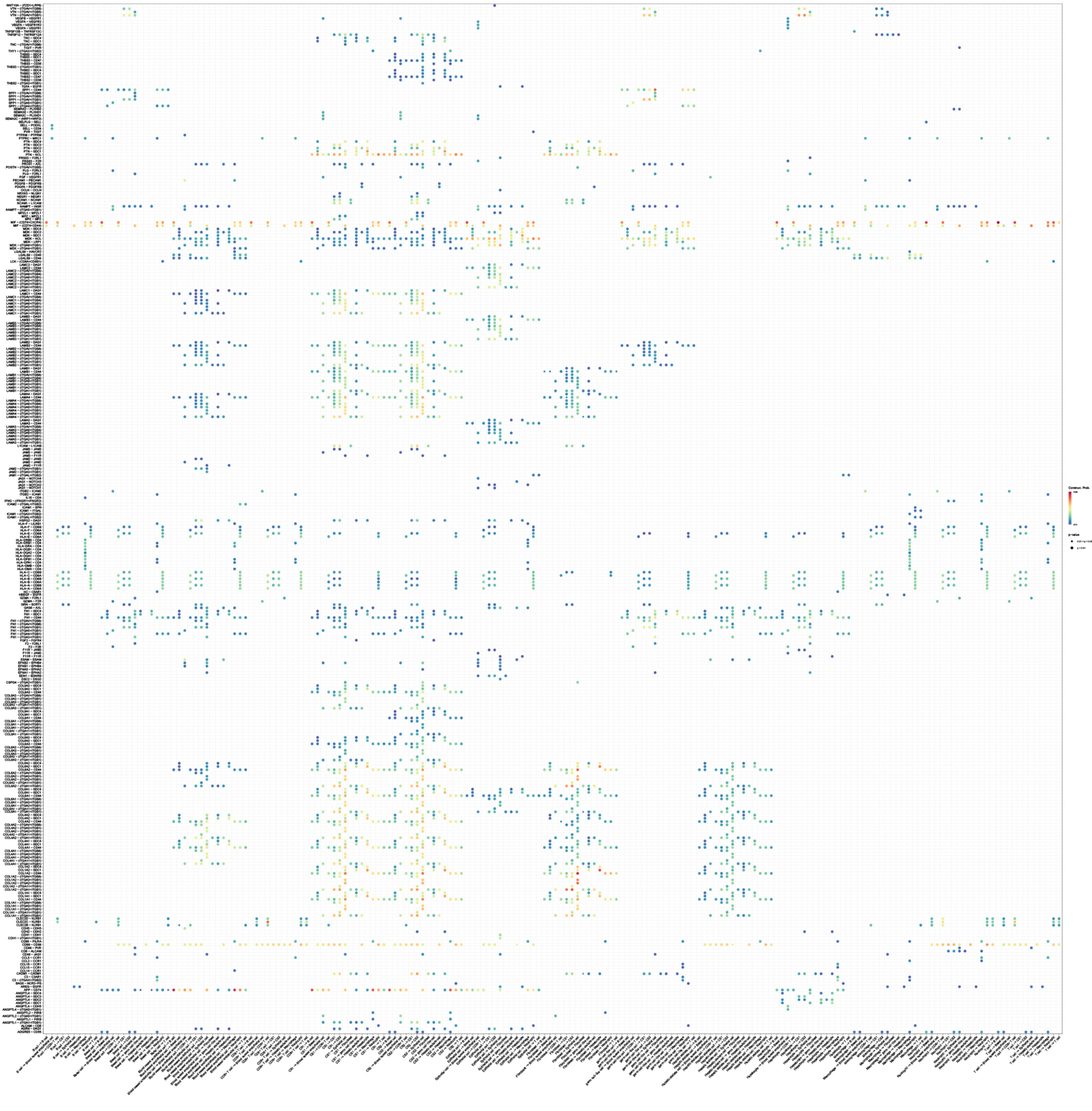

**Supplementary Figure 2**. The cell communication network composed of CS1 and CS2 subtype malignant cells and other cells in the TME.

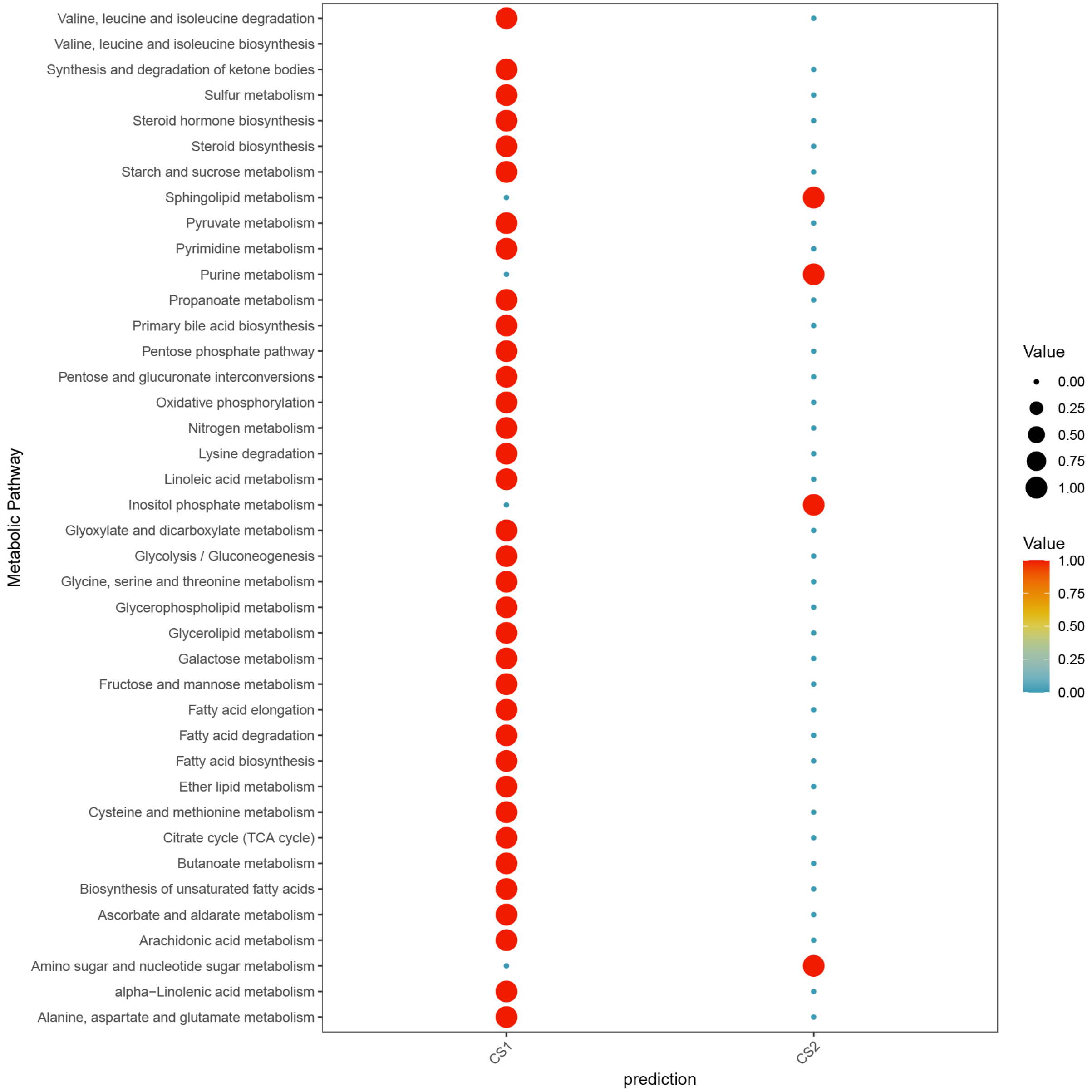


**Supplementary Figure 4.** Differences in metabolism between the two subtypes of malignant cells in GSE202642.

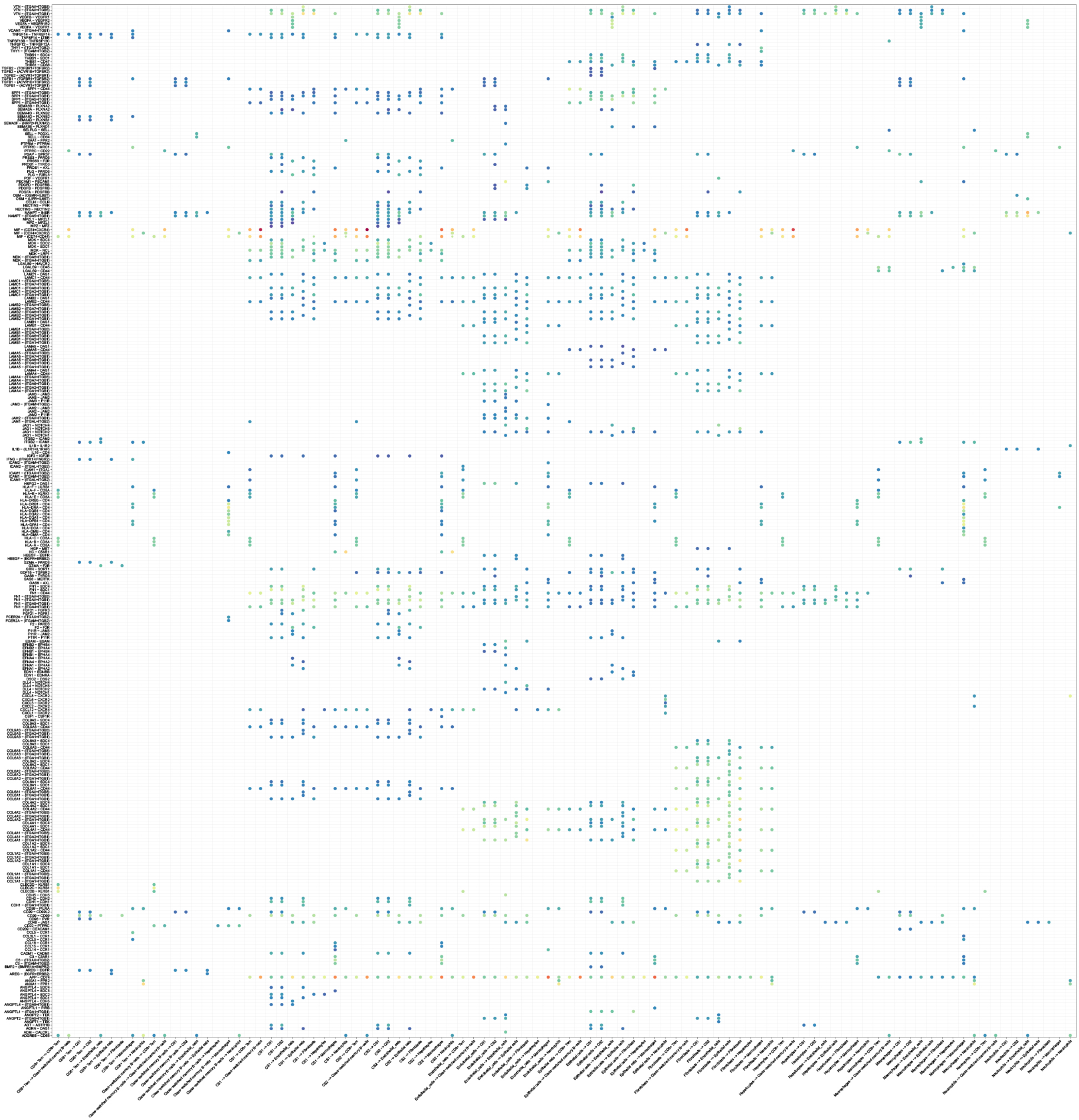

**Supplementary Figure 5**. The cell communication network composed of CS1 and CS2 subtype malignant cells and other cells in the TME.

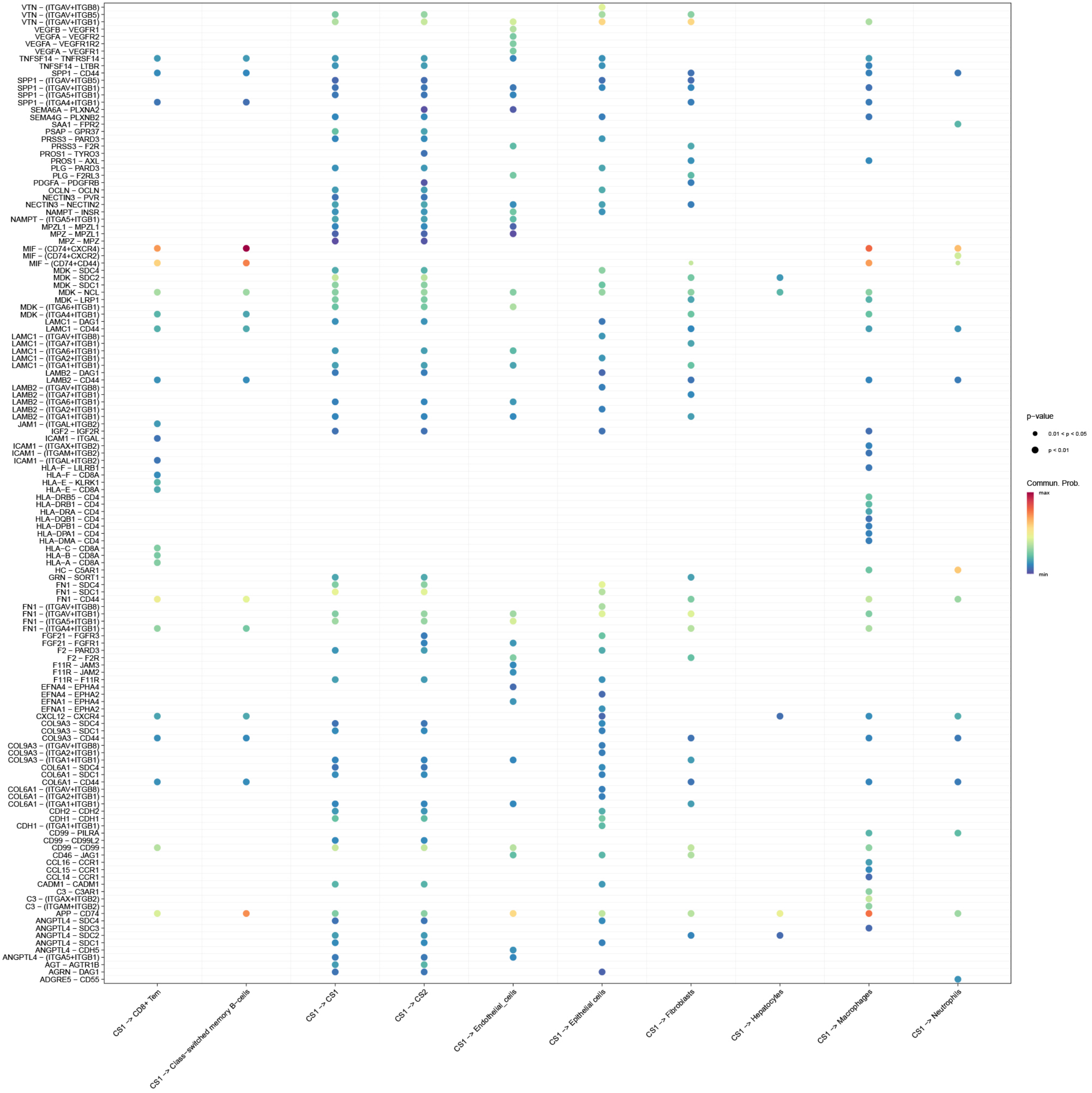


**Supplementary Figure 6**. Signaling modulation pattern of CSl subtype malignant cells on other cells in GSE202642.
